# Chill coma in the locust, *Locusta migratoria*, is initiated by spreading depolarization in the central nervous system

**DOI:** 10.1101/162198

**Authors:** R. Meldrum Robertson, Kristin E. Spong, Phinyaphat Srithiphaphirom

## Abstract

The ability of chill-sensitive insects to function at low temperatures limits their geographic ranges. They have species-specific temperatures below which movements become uncoordinated prior to entering a reversible state of neuromuscular paralysis. In spite of decades of research, which in recent years has focused on muscle function, the role of neural mechanisms in detemining chill coma is unknown. Spreading depolarization (SD) is a phenomenon that causes a shutdown of neural function in the integrating centres of the central nervous system. We investigated the role of SD in the process of entering chill coma in the locust, *Locusta migratoria*. We used thermolimit respirometry and electromyography in whole animals and extracellular and intracellular recording techniques in semi-intact preparations to characterize neural events during chilling. We show that chill-induced SD in the central nervous system is the mechanism underlying the critical thermal minimum for coordinated movement in locusts. This finding will be important for understanding how insects adapt and acclimate to changing environmental temperatures.

## Introduction

When exposed to temperatures below a critical thermal minimum (CT_min_) ^1^, chill-sensitive insects lose neuromuscular coordination and eventually enter a state of complete paralysis at the point of chill coma onset (CCO) ^2^,^3^. The susceptibility of insects to temperature extremes determines their geographical ranges and, at least for drosophilid species, CT_min_ is the best predictor of ecologically relevant cold tolerance ^4^ so understanding the mechanistic basis of CT_min_ provides insight into the evolution of ecological niche limits. For almost 50 years, research towards understanding the neuromuscular mechanisms underlying chill coma has focused on depolarization of muscle cells as the primary cause of paralysis (e.g. for recent research ^5^-^7^). Anderson and Mutchmor ^8^ came to the conclusion that neural failure is not a direct cause of chill coma in three species of cockroach because they were able to record electrical activity in the nerve cord at temperatures below the chill coma temperature. In spite of other demonstrations of chill-induced neural phenomena ^9^,^10^, the role of neural failure in determining chill coma remained unclear. Subsequent research ^11^-^13^ served only to confirm the apparent dominant role of muscle failure. However, a lack of correlation between CT_min_ and muscle depolarization has been demonstrated ^14^ suggesting that other mechanisms must be considered. Recent research in locusts ^15^ and *Drosophila* ^16^ suggests that shutdown of the central nervous system (CNS) via a spreading depolarization (SD) mechanism could have an important role to play in the entry to chill coma. The principal causes of chill coma remain to be resolved ^3^.

SD is a phenomenon which was originally described in the context of neural depression in the mammalian CNS ^17^. It plays an important role in several human pathologies ^18^ but it has also been described in insect nervous systems ^19^-^21^ where it is associated with neural depression under stressful environmental conditions ^15^,^22^. SD is characterized by a collapse of CNS ion homeostasis resulting in major changes in CNS ion concentrations, neuronal and glial depolarization and a negative DC shift in the extracellular field potential (FP) ^23^. It appears to be triggered when extracellular potassium ion concentration ([K^+^]_o_) surpasses a critical threshold and is thus dependent on activity levels in the CNS ^24^-^26^.

Our goal was to determine the relationship between behavioural chill coma and SD in the CNS of locusts. Metabolic rate drops markedly as insects enter chill coma ^27^-^29^ so we used flow-through thermolimit respirometry ^30^ as an objective measure of chill coma and related this to neuromuscular electrical activity in whole animals. Subsequent electrophysiological experiments in whole animals and semi-intact preparations showed that bursts of electrical activity in muscles during chilling have a neural origin and that SD in the thoracic ganglia indicates a point at which all neural activity in the central integrative neuropil has terminated, which would prevent coordinated movement. We conclude that SD is the mechanism for the loss of coordinated movement at CT_min_ but, because some peripheral neural activity persists after SD, which could cause uncoordinated movements, we conclude that CCO occurs at a lower temperature and is coincident with relaxation of spiracle closer muscles.

## Results

### Respirometry

The pattern of CO_2_ release from individual locusts at room temperature was variable. After an initial peak associated with handling and a build-up of CO_2_ in the chamber during the set-up, CO_2_ release could either be relatively constant or settle into a pattern of discontinuous gas exchange cycles, which have been well described ^31^,^32^. Nevertheless, during chilling, CO_2_ release dropped markedly to a level close to 0 μmoles/g/h (Fig. 1A). Subsequently there were two small peaks (Fig. 1B; < 5 μmoles/g/h) followed by a maintained period with no variation in release (1.8 ± 1.1 μmoles/g/h; n = 20), which we took to be an indication of chill coma. (Note that here and throughout the Results text, data are described as mean ± standard deviation.) Thoracic temperature at the start of experiments was 4.3 ± 0.6 °C higher than room temperature (Fig 1C; 28.8 ± 0.8 °C versus 24.5 ± 0.9 °C; paired t-test, P < 5.0 × 10^-18^; n = 20) but gradually decreased due to convective heat loss when the air flow was turned on.

**Figure 1:**
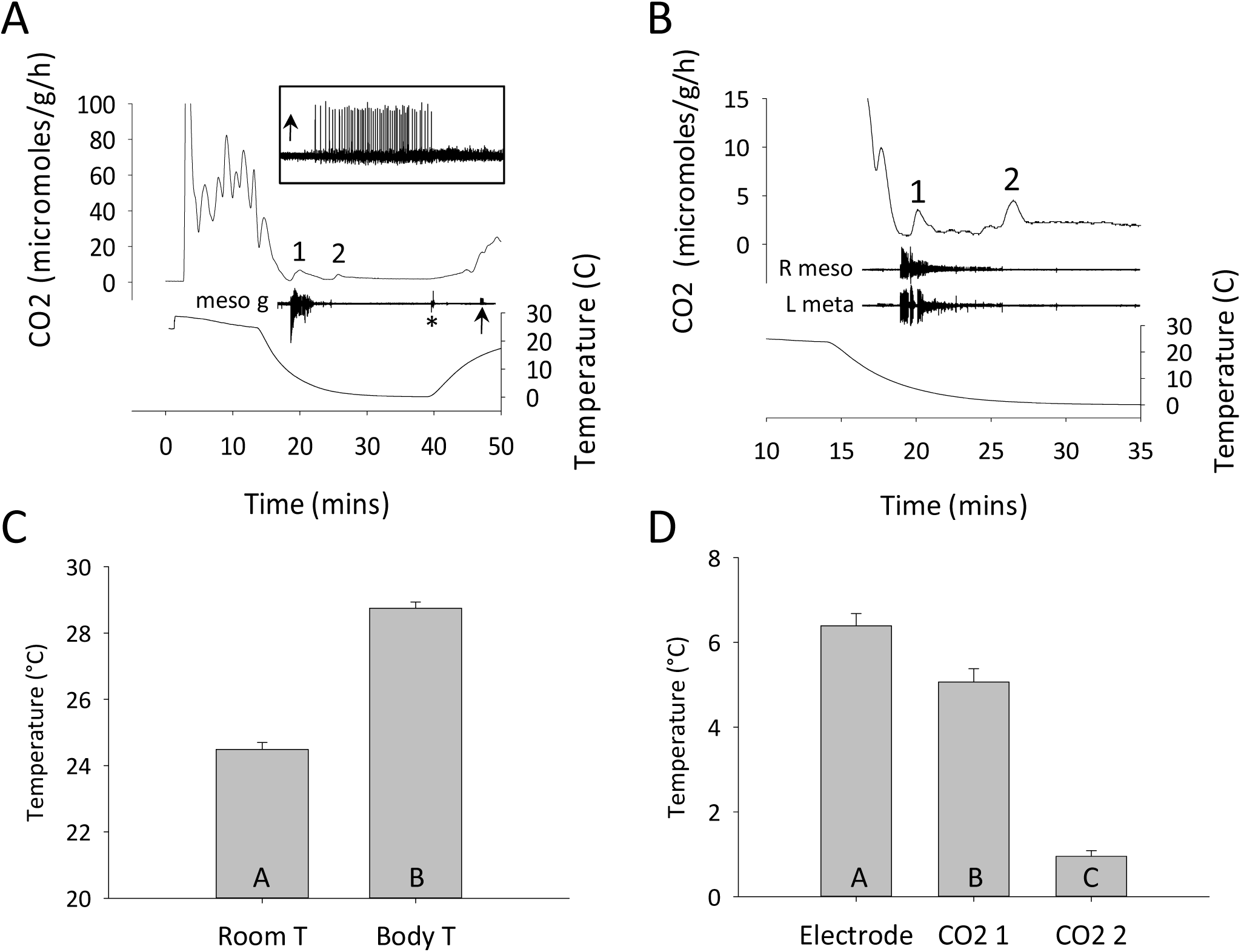
Entry into chill coma is characterized by two pulses of CO2 release associated with a burst of electrical activity in the thorax. **A.** Flow through respirometry measures of CO_2_ release time-aligned with recordings of thoracic temperature and electrical activity from an electrode inserted close to the mesothoracic ganglion (meso g). Note the step at the beginning of the temperature trace from room temperature to body temperature as the thermocouple is inserted. Asterisk marks an electrical artifact associated with removing the locust in its chamber from the chiller. 1, 2 indicate two small peaks of CO_2_ release during chilling. Arrow indicates a burst of electrical activity associated with chill coma recovery (enlarged in the inset). **B**. Similar to A, but from a different locust and focused on the two CO_2_ peaks and their relationship to electrical activity recorded electromyographically from wing muscles of the right forewing (R meso) and the left hindwing (L meta). **C**. Comparison of room temperature (Room T) and body temperature (Body T) at the start of experiments. **D**. Comparison of body temperatures at the start of electrical activity (Electrode) and at the peaks of CO_2_ release (CO2 1; CO2 2) during chilling. In C and D data are represented as mean ± standard error and different letters in the bars indicate statistically significant differences (details in Results text).

Initially we searched for a purely neural signal and were able to record bursts of electrical activity during chilling from the head capsule, a compound eye or by placing the electrode close to a thoracic ganglion (e.g. Fig. 1A). However we found that it was more reliable to record from wing muscles in the thorax (Fig. 1B). As the temperature decreased in an exponential manner there was a consistent sequence of events with the start of an intense burst of electrical activity at 6.4 ± 1.3 °C, followed by the first peak of CO_2_ release at 5.1 ± 1.4 °C and the second CO2 peak at 1.0 ± 0.6 °C (Fig. 1D; RM ANOVA P < 0.001, Holm-Sidak P < 0.001 for all comparisons; n = 20). After removal from the chiller, CO_2_ release increased gradually and then more abruptly at an inflexion point that was simultaneous with observations of the resumption of ventilatory movements and occasionally with a burst of electrical activity (Fig. 1A and inset).

### Origin of the burst of electrical activity

To determine whether the burst of electrical activity recorded in muscles originated in the muscle fibres or was dependent on activity from the CNS, we recorded from muscles during chill coma before and after lesions to the nerve supply. A problem with these recordings is that during the period immediately before CT _min_, a large proportion of the thoracic muscles, if not all of them, generate bursts of activity and the electromyographic (emg) electrode picks up distant electrical activity. To minimize the recording of distant activity we took some recordings from the extensor tibialis muscle (ETi) at a distal location in the femur on the right and left sides before and after cutting left metathoracic N5. Recordings from wing muscles and leg muscles revealed the early burst noted in the previous section and a later one that was more prominent in leg recordings (Fig. 2A). In this dataset from 38 locusts, thoracic temperature at the start of the experiment was 3.6 ± 1.4 °C higher than room temperature, the start of the first muscle burst occurred at a thoracic temperature of 6.2 ± 1.8 °C and the start of the second muscle burst started at a thoracic temperature of 1.0 ± 0.7 °C.

**Figure 2:**
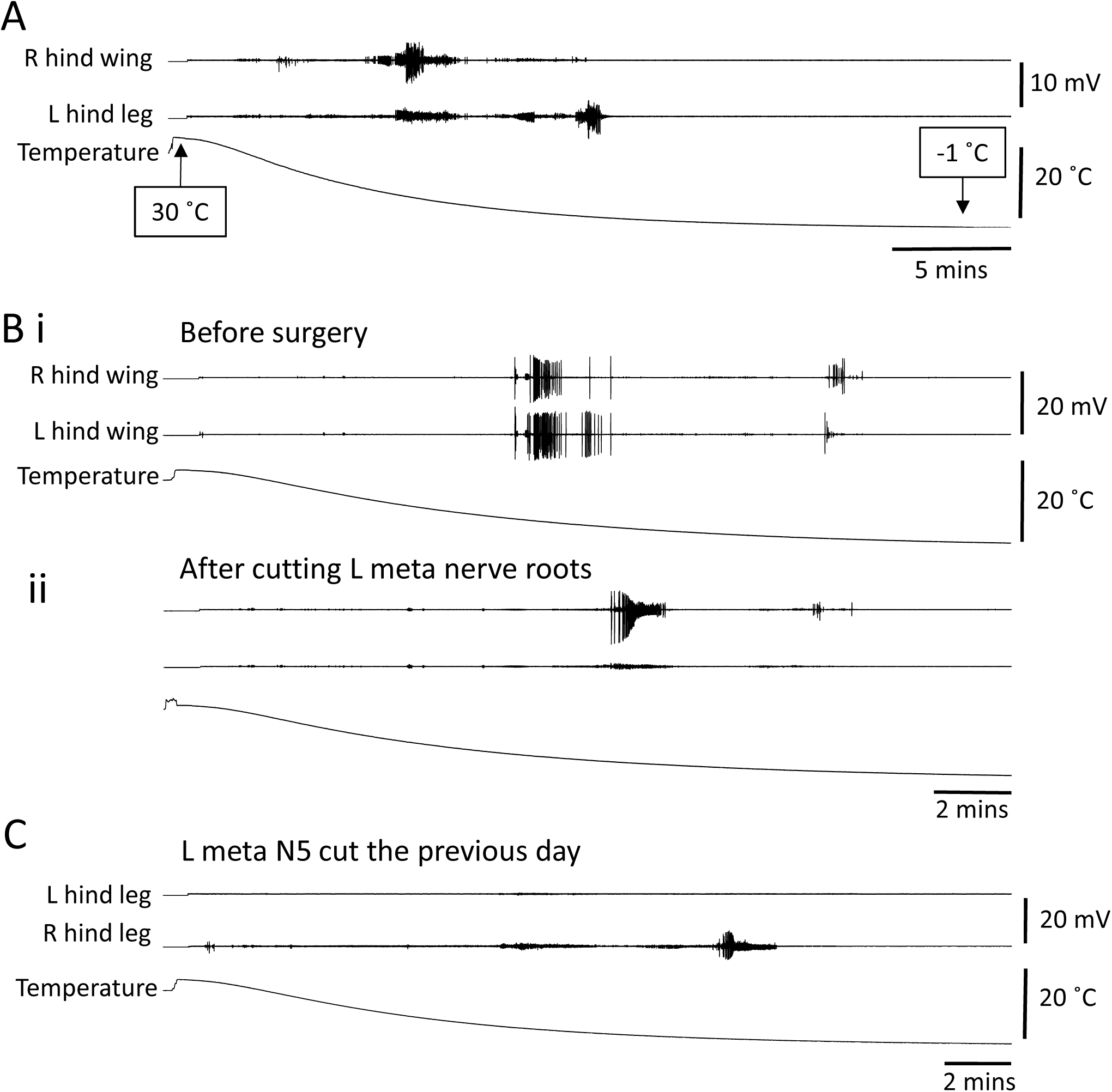
Chilling causes bursts of electrical activity in wing and hind leg muscles that originate in the CNS. **A.** Emg recording from a right hind wing muscle and a left hind leg muscle during chilling from thoracic temperature of 30 °C to −1 °C. Note bursts of muscle activity during chilling. **B**. Similar to A, but recording is from right and left hind wing muscles. **i**. Before surgery. **ii**. After cutting the nerve root supplying the left hind wing muscle. Note that the surgery eradicates large amplitude bursts of muscle activity during chilling (amplitude scales same as in i). **C**. Similar to A and B, but recording is from left and right hind leg extensor tibialis muscles in a locust that had the nerve root supplying the left muscle cut the previous day. Note the lack of any muscle bursts associated with chilling in the left muscle.

For 16 locusts during entry into chill coma we recorded from thoracic flight muscles on the right and left, before and after cutting all the nerve roots on the left side of the metathoracic ganglion. Large amplitude bursts of electrical activity which could be unequivocally attributed to the implanted muscle were eradicated (Fig. 2B). The same result was obtained after complete removal of the metathoracic ganglion in 8 locusts (not shown). In 12 locusts, cutting metathoracic N5 eradicated or prevented the recording of any large amplitude bursts of electrical activity in ETi during entry into chill coma (Fig. 2C).

Large amplitude bursts of electrical activity in muscles recorded during entry into chill coma were undoubtedly a consequence of muscle fibre excitability. Nevertheless, their occurrence was dependent on an intact nerve supply indicating that they were triggered by events originating in the CNS.

### Spreading depolarization in the CNS

SD in the thoracic ganglia was characterized by an abrupt surge of [K^+^]o in the neuropil (Fig. 3A; peak = 112 ± 59 mM from a baseline = 7.2 ± 3.0 mM; n = 6). The ionic disruption was monitored in other experiments by recording the field potential (Fig. 3B), which shows an abrupt negative DC shift, which is coincident with the [K^+^]o surge (Fig. 3C; negative peak = −38.4 ± 9.5 mV; n = 12). SD was preceded by and followed by positive peaks of FP (Fig. 3C; preceding peak = 7.4 ± 1.9 mV; following peak = 5.7 ± 2.9 mV; n = 12). Recording at two different locations in the mesothoracic ganglion showed that the event occurred at different times suggesting that the disturbance spread throughout the neuropil (Fig. 3D). The median latency between the events at different locations was 13 s (IQR 6 - 14 s; n = 7), which, assuming an electrode spacing of ~0.4 mm and spread from one location to the other, gives an estimate speed of propagation of 1.8 mm/min (IQR 1.7 - 4 s; n = 7). Simultaneous recording of FP in the metathoracic ganglion and extracellular neural activity from metathoracic N5 (cut distally), which supplies the ETi and other muscles of the hindleg, showed a burst of action potentials prior to the chill-induced SD and another burst coincident with recovery of the FP after chilling (Fig. 3E, representative of n = 14). It should be noted that, in contrast to the recordings from the neuropil (see below), in about half of the nerve root recordings, action potentials were evident after the negative DC shift (Fig. 3F).

**Figure 3:**
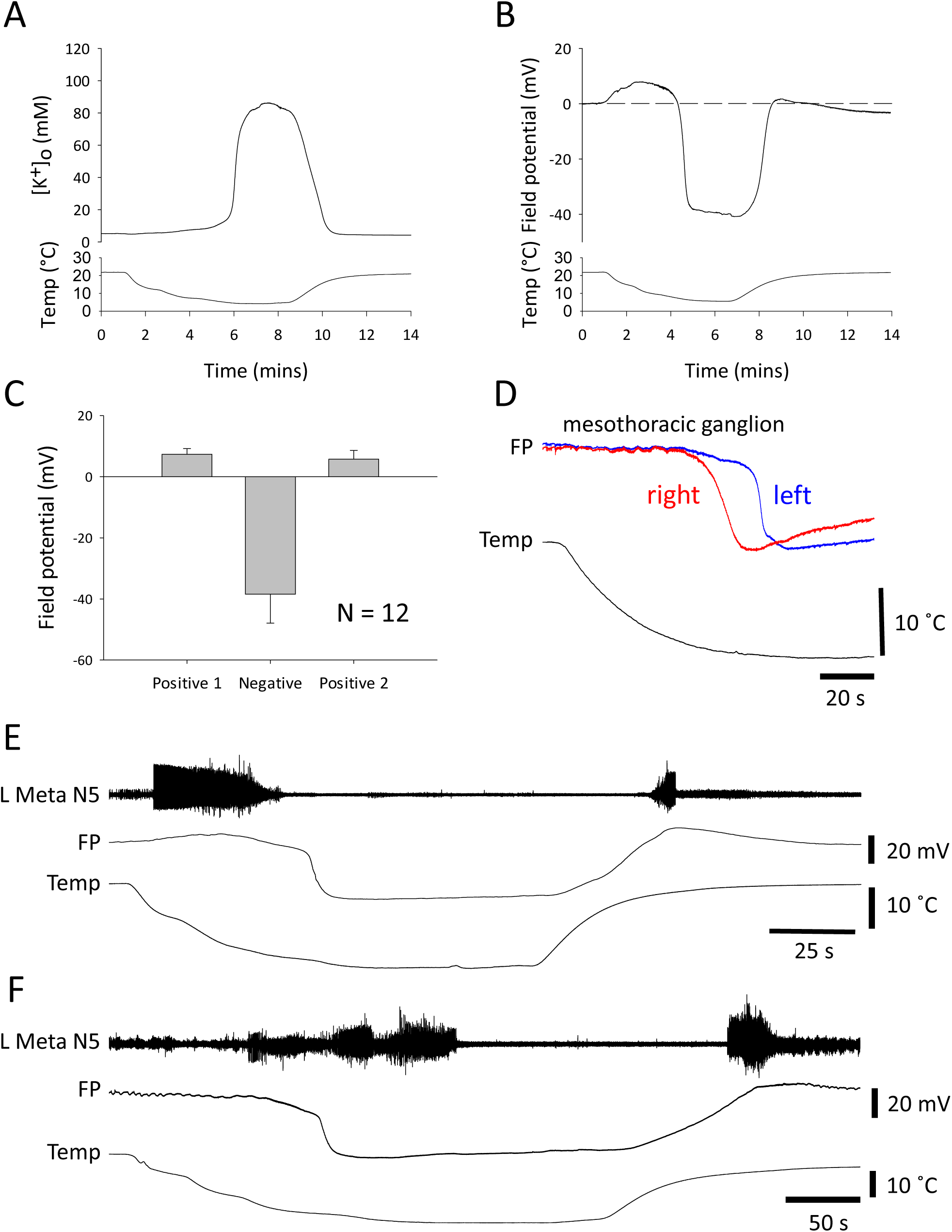
Chilling causes SD in the CNS characterized by a negative shift of extracellular field potential. **A.** Representative example of the surge of extracellular potassium [K^+^]o associated with chilling. **B**. Representative example of a field potential (FP) recording from the neuropil of the metathoracic ganglion. The surge in [K^+^]o and the abrupt negative shift in FP occur simultaneously and are hallmarks of SD. **C**. Quantification of the three phases of FP changes during chill-induced SD (positive 1, negative, positive 2). Data represented as mean ± standard deviation. **D**. Simultaneous recordings of FP from two locations in the mesothoracic ganglion during chilling. The traces have been scaled to be the same amplitude for display purposes. Note the time lag between SD recorded in the right and left electrodes. **E**. Representative example of simultaneous recording of nerve activity in the left metathoracic nerve 5 and FP in the metathoracic ganglion during chilling. Note the bursts of neuronal activity prior to SD and on recovery. **F**. Same as E, but in a different preparation to show that peripheral electrical activity could be recorded after SD during chilling.

### Axonal performance of the DCMD neuron

To characterize the effects of chilling on an important, identified interneuron we recorded extracellularly from the axon of the descending contralateral movement detector (DCMD) interneuron in the connective between the pro- and mesothoracic ganglion during continuous visual stimulation. Immediately the saline temperature started to drop, the amplitude of the extracellular, triphasic action potential began to decrease and this continued until action potential conduction failed (Fig. 4A). Axonal performance during chilling was assessed using two suction electrodes to measure conduction delay at different firing frequencies (Fig. 4B). Associated with its functional role in predator avoidance, the DCMD axon is adapted to conduct large action potentials at high firing frequencies with minimal drop in conduction velocity ^33^,^34^. As temperature decreased, the axonal performance was increasingly impaired, showing a reduction in high frequency firing and an inability to maintain high conduction velocity (short anterior to posterior latency). DCMD conduction failed at higher temperatures than SD in the metathoracic ganglion (Fig. 4C; DCMD fail – 9.9 ± 2.6 °C; FP drop – 7.7 ± 2.6 °C; DCMD recovery – 17.3 ± 2.1 °C; RM ANOVA P < 0.001; Holm-Sidak P < 0.004 for all pair-wise comparisons; n = 18).

**Figure 4:**
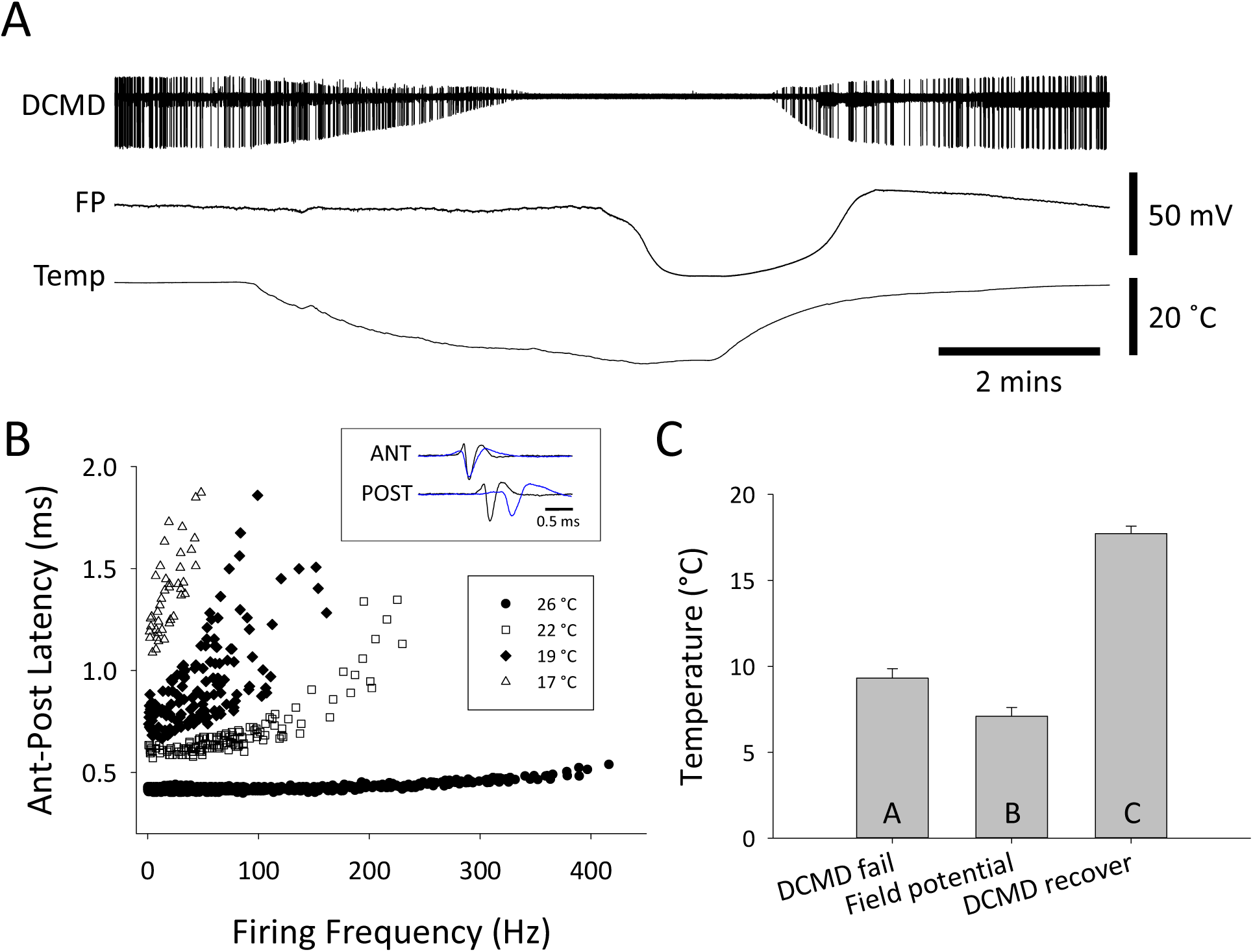
Chilling impairs performance of the DCMD neuron prior to SD in the mesothoracic ganglion. **A.** Representative example of simultaneous extracellular recording in a semi-intact preparation of DCMD action potentials in the pro- mesothoracic connective and FP in the metathoracic ganglion during chilling. DCMD was continuously visually stimulated. Note failure and recovery of action potential conduction in the DCMD. **B**. Representative example of loss of DCMD axonal performance during chilling. Performance is represented by the ability to conduct action potentials at high frequency with minimal decrease in conduction velocity, represented here as the latency to conduct an action between an anterior and a posterior electrode (Ant-Post Latency). The inset shows an overlay of recordings at room temperature (26 °C) and at 19 °C, aligned on the triphasic extracellular action potential in the anterior electrode (ANT). **C**. Comparison of saline temperature at failure of axonal conduction in DCMD (DCMD fail), the negative DC shift of FP (Field potential) and recovery of axonal conduction in DCMD. Data are represented as mean ± standard error and different letters in the bars indicate statistically significant differences (details in Results text).

### Intracellular activity during chilling

Wing muscle motoneurons in the mesothoracic ganglion were identified by activating the flight central pattern generator with wind stimulation of the head (Fig. 5). Elevator motoneurons fired bursts of action potentials in antiphase with activity of the dorsal longitudinal (DL) wing depressor muscle (Fig. 5B). Motoneurons started to depolarize as soon as the temperature dropped and then generated a burst of action potentials with steadily decreasing amplitudes until action potential generation failed. Chilling also induced a barrage of postsynaptic potentials (Fig. 5C), which contributed to firing of the motoneurons, indicating that premotor interneurons were also firing action potentials during chilling. These postsynaptic potentials were not evident during the chill-induced neuronal depolarization. On return to room temperature, some motoneurons generated a burst of action potentials with steadily increasing amplitude as their membrane potentials repolarized (Fig. 5A), others simply repolarized (e.g. Fig. 6B).

**Figure 5:**
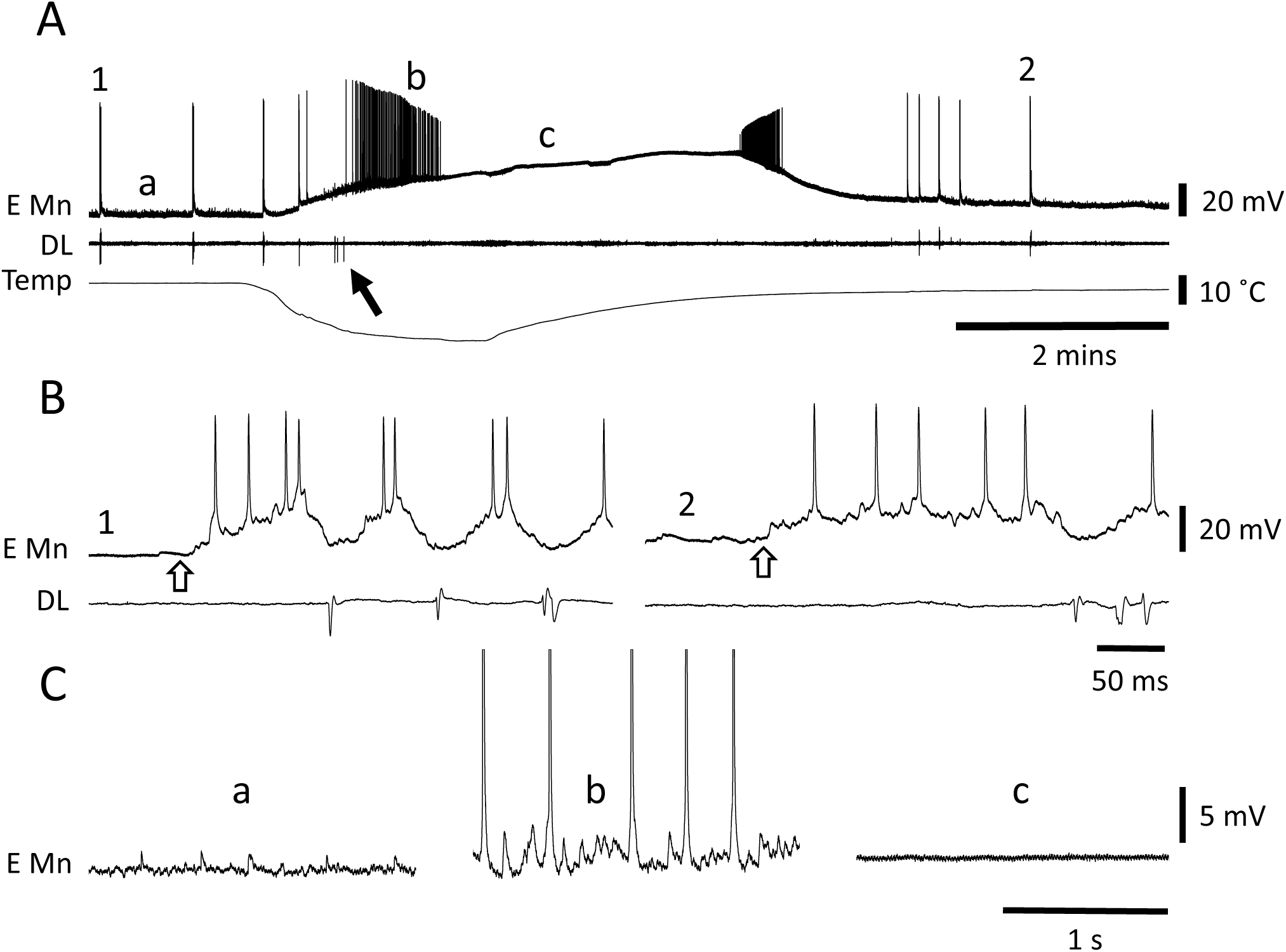
Wing muscle motoneurons depolarize and are inactivated during chilling. **A.** Representative intracellular recording from a wing elevator muscle motoneuron (E Mn) and emg recording from a dorsal longitudinal depressor muscle (DL) during failure and recovery associated with chilling and return to room temperature. 1 and 2 marks portions of the traces expanded in B. a, b and c mark portions of the traces expanded in C. The arrow points to a DL burst associated with chilling. **B**. During expression of the centrally generated flight motor pattern induced by wind stimulation of the head (start indicated at open arrow) the intracellular recording could be identified as an elevator motoneuron due to the phase of its activity relative to DL. 1– prior to chilling. 2 – after recovery. **C**. Illustration of subthreshold synaptic activity in the elevator motoneuron. **a** – background synaptic activity before chilling. **b** – high frequency synaptic activity during chilling illustrating firing of premotor interneurons. Note that action potentials have been clipped. **c** – no synaptic activity during chill-induced motoneuron depolarization.

**Figure 6:**
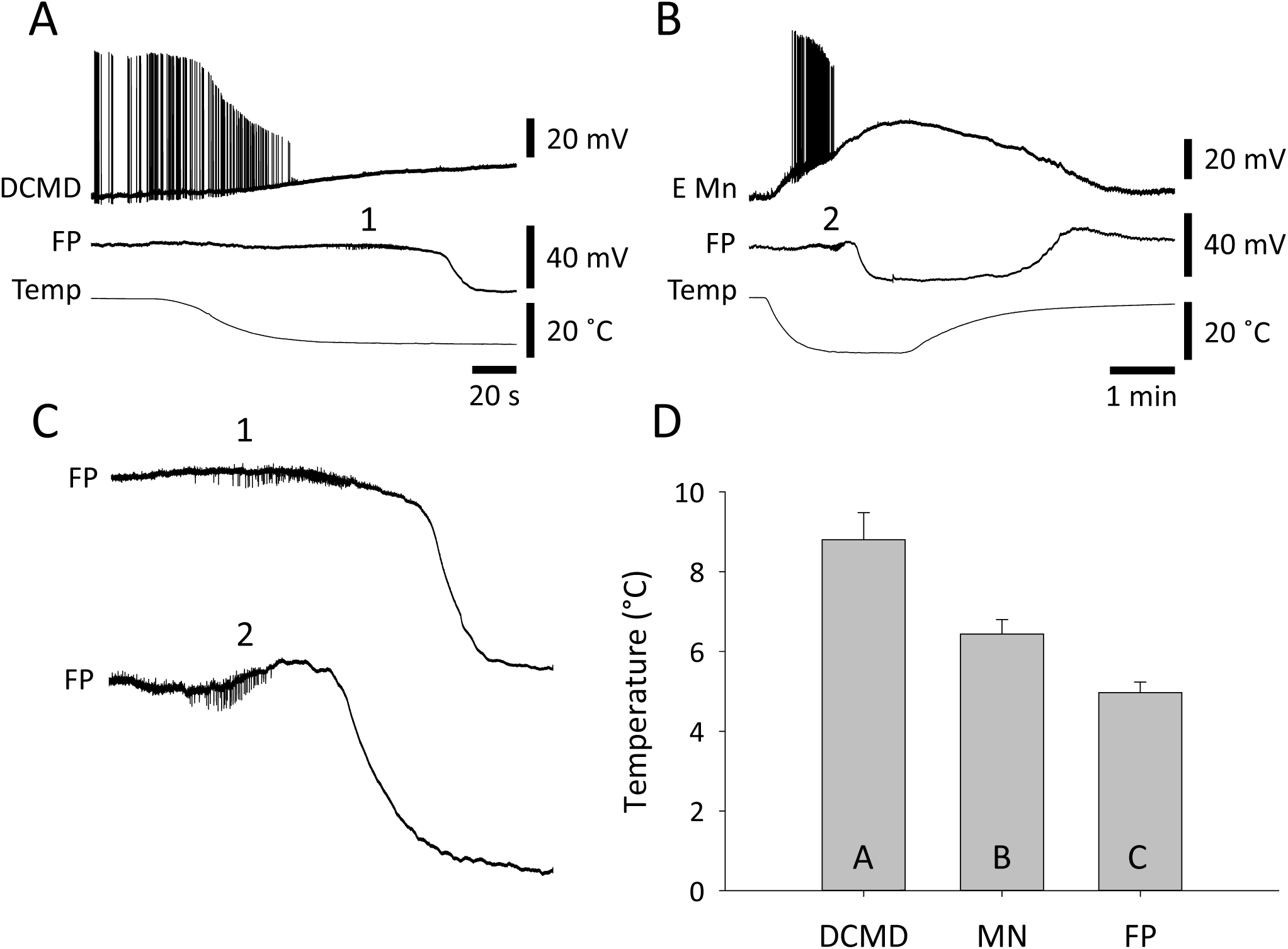
SD in the thoracic ganglia occurs after neurons in the neuropil are depolarized and have ceased firing. **A.** Simultaneous intracellular recording from the DCMD axon and FP in the metathoracic ganglion during chilling. 1 indicates a portion of the FP trace expanded in C. Note that SD (negative DC shift of FP) occurs after depolarization and failure of DCMD. **B**. Simultaneous intracellular recording from an elevator motoneuron (E Mn) and FP in the mesothoracic ganglion during chilling. 2 indicates a portion of the FP trace expanded in C. Note that SD occurs after depolarization and failure of E Mn. **C**. Enlargement of the FP traces in A and B shows extracellular pick up of bursts of neuronal activity (at 1 and 2) that cease prior to SD. **D**. Comparison of saline temperatures at failure of DCMD, failure of wing muscle motoneurons (MN) and FP. Data are represented as mean ± standard error and different letters in the bars indicate statistically significant differences (details in Results text).

Simultaneous FP recording with intracellular recording from the axon of the DCMD (Fig. 6A) or wing muscle motoneurons (Fig. 6B) showed action potential failure prior to SD. Often the FP electrode recorded extracellular single unit activity of interneurons which generated bursts that stopped prior to SD (Fig. 6C). Action potential generation failed first in the DCMD axon (8.8 ± 2.9 °C; n = 18) and then in wing muscle motoneurons (6.4 ± 1.6 °C; n = 19) prior to SD (5.0 ± 1.3 °C; n = 26) (Fig. 6D; ANOVA P < 0.001; Holm-Sidak pair-wise comparisons: DCMD vs. FP P < 0.001, DCMD vs. MN P = 0.001, MN vs. FP P = 0.016). Chill-induced depolarization in the best recordings was 19.4 ± 5.4 mV (n = 5) in the DCMD axon and 28.0 ± 8.1 mV (n = 7) in wing muscle motoneurons. Given resting membrane potentials around −60 mV, these neurons did not depolarize completely during chill-induced SD.

It is important to note that, in contrast with the nerve root recordings (metathoracic N5), we never observed any indication of neural activity in the neuropil after chill-induced SD.

## Discussion

We found that in locusts, as in other chill-sensitive insects ^11^-^13^, there was a characteristic burst of action potentials in wing muscles during chilling. This was evident both in the emg recording from wing muscles of intact locusts and in the intracellular recordings from wing muscle motoneurons in semi-intact preparations. We were thus able to correlate neural events in the CNS, such as neuronal membrane potential and extracellular FP, with entry to chill coma of intact locusts. Our primary conclusions are: 1. that the electrical activity in muscles associated with CT _min_ and chill coma required an intact efferent innervation; and 2. that SD in the thoracic neuropil was the point at which coordinated motor activity was effectively precluded i.e. that chill-induced SD is the mechanism underlying the loss of coordinated movement at CT_min_.

We were expecting to see an abrupt drop in metabolic rate, evident as a decrease in CO_2_ release, as locusts entered chill coma. Thus we were surprised to record consistently two small pulses of CO_2_ release after a major drop during chilling. The first of these occurred at a slightly lower temperature than bursts of electrical activity in the thorax. All the recordings indicate that many motoneurons and interneurons were active at high frequencies at about the same time and this would have caused major ionic disturbances, activating the Na^+^/K^+^- ATPase and increasing metabolic rate. Therefore we interpret the first CO_2_ pulse as an indication of metabolic rate increase associated with chill-induced neuronal hyperactivity. The second CO_2_ pulse occurred at a significantly lower temperature, lower than the temperature when the Na^+^/K^+^-ATPase was expected to have stopped operation (= SD, see below), and immediately prior to CCO, indicated by no further electrical activity and a flat line in the CO_2_ recording. In the absence of muscle tone, spiracles are open such that during chill coma there is a continuous slow release of CO_2_ ^27^,^35^. In addition, *Tenebrio molitor* beetles show a prominent spike in CO_2_ release as they enter chill coma, which is interpreted as being caused by spiracle opening ^29^. Thus we suggest that the second CO2 pulse was caused by relaxation of spiracle closer muscles at CCO and release of tracheal contents.

There are many approaches for estimating critical thermal limits, including acute exposures and ramped temperatures, and they have their advantages and disadvantages ^30^. Because we were less concerned with determining environmental temperature limits than the core body temperature at particular neural events, we chose treatments that had comparable effects on thoracic temperature in intact animals and semi-intact preparations. In intact locusts, acute exposure to a low temperature (-1 °C) caused an exponential drop in thoracic temperature that took ~10 mins to plateau. In semi-intact preparations, the ramped chilling of the saline inflow also caused an exponential drop in thoracic temperature that took 5-8 mins to plateau. The rapid drop in temperature was appropriate for intracellular recording from neuronal dendrites, which, for technical reasons, may not be long-lasting. However, we recognize that locusts are unlikely to experience such rapid temperature changes in the wild. Using a more ecologically relevant, slow temperature change could affect the absolute values of CT_min_ and CCO but would be unlikely to change their underlying mechanisms. It is possible that very slow depolarization of neurons would minimize or prevent action potential bursts due to neuronal accommodation ^36^, or due to compensatory membrane or ionic adjustments that minimize or prevent depolarization. Nevertheless, at some point CT_min_ will be reached and the locust will enter a coma. We predict that CT_min_ at the end of a slow change in ambient temperature will be marked by SD in the CNS.

Our different methods make it difficult to compare our measures with previous descriptions of CT_min_ in *Locusta migratoria*. This has been reported around 0 °C using slow ambient temperature ramps and noting the point at which no more activity is observed ^5^. In that investigation changing the cooling rate between 0.02 and 0.18 °C/min didn’t have a marked effect on CT_min_ but the cooling rate in our experiments was around an order of magnitude faster than their fastest rate. Also we found that, in still air, thoracic temperature was maintained around 4 °C higher than ambient room temperature. Thus, if CT_min_ is defined as an exposure temperature for loss of coordination, our measures of thoracic temperature at that point would be expected to be higher than CT_min_. Moreover, the temperature when no more activity is observed, as opposed to a loss of coordination, might be more appropriately referred to as CCO.

The chill-induced SD that we characterize here has interesting features suggesting we should re-evaluate our model for their generation ^37^. In that model the abrupt onset of SD is proposed as resulting from a positive feedback cycle involving neuronal depolarization and firing, which increases [K^+^]o, which causes depolarization. At low temperatures, however, it is clear that neuronal depolarization and firing occurs prior to and is distinct from the onset of SD, which occurs after neuronal activity in the neuropil has been silenced. One possibility is that SD onset is primarily a glial event, perhaps associated with an abrupt failure of glial spatial buffering of [K^+^]o via gap junctions ^38^. This would be counter to the current orthodoxy regarding SD in vertebrates ^18^. Alternatively, the Na^+^/K^+^-ATPase clearly has an important role to play ^21^,^22^ and in mammalian brains it has been suggested that, at a critical threshold, the Na^+^/K^+^-ATPase is converted into a large conductance open cationic channel that provides the ion current underlying SD ^39^. This intriguing possibility remains to be exa_min_ed but the notion that SD onset represents a well-defined point at which the ion pump fails is consistent with our data. Prior to SD the ion disturbance caused by chill-induced neuronal hyperactivity is adequately compensated by hard-working, energy-consuming pumps that transiently increase metabolic rate but maintain a low [K^+^]o. It is unclear what triggers pump failure at low temperatures but it is more than just a thermodynamic slowing of operation.

Defining CT_min_ and CCO in insects has been problematic but a consensus has emerged that distinguishes between general impairment of function, loss of coordination (at CT_min_) and complete absence of electrophysiological activity and movement (at CCO) as temperature drops ^1^-^3^. This sequence is clearly evident in our recordings. Immediately temperature starts to drop (thoracic temperature: 30 - 10 °C) neuronal and circuit properties change in predictable ways ^40^,^41^ and behaviour is slowed but not impaired. Then (10 - 6 °C) bursts of neuronal activity occur (see also ^10^) and neurons depolarize and are incapacitated at different times. This is a period when behaviour is impaired but still possible. For example, conduction failure of DCMD will impair rapid predator evasion and depolarization-induced silencing of motoneurons will prevent activation of muscles. SD in the CNS (5 °C) occurs after all evidence of neuronal communication (action potentials and synaptic potentials) in the neuropil has vanished and it represents a well-defined point after which no neural integration is possible (i.e. loss of coordinated movement at CT_min_). SD is a phenomenon that is confined to neuropil in mammals and insects ^15^,^19^ and it does not propagate into nerve roots or connectives. Thus, after SD it is still possible to record peripheral electrical activity in muscles and nerve roots (5 - 1 °C) but this is uncoordinated and comes to an end when membrane excitability in nerve roots and muscles completely fails (i.e. absence of electrophysiological activity and movement at CCO).

At the same time as this sequence of neural events during chilling, there are well-described changes in muscle properties ^5^,^6^,^42^ that contribute to behavioural impairment and paralysis. The relative importance of nerve and muscle for chill-induced neuromuscular paralysis is likely to vary in different species and under different conditions. However, crossing a threshold of muscle fibre depolarization during chilling does not correlate completely with chill coma temperatures in *Drosophila* species that vary in cold tolerance ^14^. To understand the mechanisms involved in acclimation and adaptation to cold environments, it will be necessary to determine the role of chill-induced SD.

## Author Contributions

Conceptualization: RMR. Methodology: RMR, KES. Investigation: RMR, KES, PS. Resources: RMR. Writing – original draft preparation: RMR. Writing – review and editing: RMR, KES, PS. Visualization: RMR. Supervision, project administration and funding acquisition: RMR.

## Acknowledgements

We thank JD Gantz for help collecting some of the data and for comments on a previous version of the manuscript. Funded by the Natural Sciences and Engineering Research Council of Canada. The authors declare no conflicts of interest.

## Data Availability

Data available on request by contacting RMR.

## Methods

### Animals

Locusts (*Locusta migratoria*) in the gregarious phase were obtained from a crowded colony maintained in the Biosciences Complex at Queen’s University. We used mature adults aged 3-6 weeks past the final moult. To avoid potential variance in the data associated with female reproductive status, we used only males. This may affect the absolute value of some of the reported measures but is unlikely to affect the general findings. The colony was kept with a room temperature of 25 °C and a photoperiod of 12:12 (lights on at 7:00 a.m.). During the light period, incandescent light bulbs in the cages increased temperature to ~30 °C and the locusts were able to thermoregulate behaviourally to preferred body temperatures. They were fed once a day with wheat grass and a mixture of bran, dried milk powder and yeast. Locusts selected for experiments were taken at random from the colony and held in the laboratory in ventilated plastic containers.

### Dissected preparations

Semi-intact preparations to reveal the thoracic nervous system were made by removing the wings and legs of a locust, pinning it to a cork board and dissecting from a dorsal approach ^43^. The gut was cut posteriorly and pulled out of the cavity. Overlying tissue was removed and the meso- and metathoracic ganglia were raised and stabilized on a metal plate. Nerve roots 3 and 4 of both sides of the two ganglia were cut to minimize movements associated with activation of thoracic muscles. Nerves 1 were not cut to preserve the innervation of the dorsal longitudinal (depressor) wing muscles. Nerves 5 were cut when the legs were removed and in some experiments nerve 5 on the left side of the metathoracic ganglion was cut closer to the ganglion to ensure that only efferent neural activity was recorded. The thoracic cavity was perfused with standard locust saline (in mM: 147 NaCl, 10 KCl, 4 CaCl _2_, 3 NaOH and 10 HEPES buffer; pH = 7.2; chemicals from Sigma-Aldrich) using either a peristalic pump (Peri-Star, WPI Inc.) or gravity flow from a reservoir suspended above the preparation.

### Temperature measurement and control

For experiments using intact animals, an IT-18 copper/constantan thermocouple (0.064 mm diameter; time constant 0.1 s) connected to a BAT12 thermometer (Physitemp Instruments Inc.) was inserted into the thoracic cavity through a hole in the arthrodial membrane under the anterior edge of the pronotum to a distance of ~1 cm. Locusts were lightly restrained on a narrow (0.5 X 4 cm) cork platform and held in the body of a 10 ml plastic syringe. For the respirometry experiments the syringe was sealed, with tubing for air to be pumped through the chamber and then placed in a chiller (Julabo F10) containing ethylene glycol held at −1 °C. The thermocouple wire and any copper wire electrodes were compressed between the stopper with the inlet tube and the body of the syringe to prevent air escaping. The assembly was inserted vertically into the chiller fluid with the region containing the locust submerged and the inlet region, with the wires, held out of the fluid. For whole animal electromyography without respirometry, the locusts were implanted with electrodes and thermocouple, restrained and held in the body of a 10 ml syringe that was cut and completely open at both ends. This assembly was then placed horizontally into a 500 ml beaker that was submerged to a depth of about 3 inches in the chiller held at −3 °C.

For experiments using semi-intact preparations the thermocouple was placed adjacent to the mesothoracic ganglion. The saline was chilled by placing different lengths of the saline inlet tubing in a slurry of water, ice and salt (−7 to −10 °C) and controlling the flow rate so that it was fast enough to avoid both freezing in the slurry and warming between the slurry and the preparation, and slow enough to cool down during its passage through the slurry. Using this method it was possible to reduce the temperature of the saline in the thoracic cavity to around −1 °C.

### Flow-through respirometry

We measured CO_2_ release using flow-through respirometry. Pre-weighed, whole animals implanted with a thermocouple and one or two copper wire electrodes were sealed in a 10 ml syringe as described above. Air was pumped from a reservoir filled from a standard air supply (i.e. not from the room air, which could have a variable gas composition), through the syringe and over the locust at 100 ml/min before being dehydrated and sent through a CO_2_ analyzer (Qubit Systems S151). CO_2_ concentration was measured every second, digitized (Vernier Labpro) and displayed using Loggerpro software. Electrode and temperature recordings were digitized and recorded separately, as described below, and synchronized with the CO_2_ measurement in the subsequent analysis by taking into account the flow lag time from the locust to the analyzer. CO_2_ concentration was converted into a mass-specific rate of CO_2_ release in μmoles/g/h.

Electrodes were implanted into a locust before it was placed in the syringe. Then the thermocouple was inserted to give a measurement of room temperature followed by thoracic temperature. Only then was the syringe sealed and the air flow turned on, which meant that CO2 released from the locust while the thermocouple was being inserted collected in the syringe and was measured as a peak of CO_2_ concentration when the air flow reached the analyzer (allowing a measurement of the flow lag time). CO_2_ release was recorded for 15 mins before the assembly was placed in the chiller and for 15 mins after an indication the locust had entered chill coma before being taken out of the chiller.

### Electrophysiology

Extracellular recordings from whole animals were made by inserting copper wire electrodes (50 μm diameter, insulated except at the cut end) ~0.5 mm through small pinholes made in the cuticle and securing them with a small drop of molten wax. Signals from monopolar electrodes were amplified (Grass P15, single-ended configuration, low frequency cut off at 3 Hz and high frequency cut off at 3 kHz), digitized (1 kHz sampling rate, MiniDigi 1A, Molecular Devices) and recorded using Axoscope 10.3 software (Molecular Devices). The locust was grounded with a separate chlorided silver wire connected to the ground terminal of the P15 amplifier and inserted into the thorax under the anterior edge of the pronotum. The same electrodes and amplifier were used to record flight motor patterns from the dorsal longitudinal (DL) wing depressor muscle in semi-intact preparations.

In semi-intact preparations, extracellular nerve recordings were made using suction electrodes (A-M Systems) with custom-made tips pulled from glass capillaries. Signals were amplified with a differential AC amplifier (A-M Systems model 1700, low cut off at 1 Hz and high cut off at 10 kHz). The preparation was grounded with a chlorided silver wire inserted into the locust’s abdomen. Intracellular recordings were made with glass microelectrodes (20-40 megohm, back-filled with 3 M KCl) and amplified with a model 1600 Neuroprobe amplifier (A-M Systems). K^+^-sensitive electrodes were made using silanized glass capillaries, pulled to 5-7 megohm, filled at the tips with Potassium Ionophore I-Cocktail B (5% valinomycin; Sigma-Aldrich) and back-filled with 500 mM KCl ^19^. Voltage from the K^+^-sensitive electrode was referenced against voltage recorded with an extracellular glass microelectrode (5-7 megohm; back-filled with 3 M KCl) positioned just adjacent and amplified with a DUO773 amplifier (WPI Inc.). The electrodes for [K^+^]o measurement were calibrated at room temperature using 15 mM and 150 mM KCl solutions to ensure that the electrode sensitivity fell between a range of 54 to 58 mV for a 10 fold change in [K^+^]. Field potential recordings were made with the same electrodes used as the reference for [K^+^]o measurement. All signals from semi-intact preparations, including the signal from the thermocouple, were digitized using a 1440A digitizer (100 kHz sampling rate; Molecular Devices) and recorded using Axoscope 10.3 software.

### Surgery

For some whole animal experiments recordings were made during entry to chill coma before and after, or only after, lesioning the nervous system. In most cases the surgery could be completed within 5minutes, while the locust was still in chill coma. A small flap (0.5 mm square) was cut into the midline metathoracic sternal cuticle, just posterior to the second spina and the junction with the mesothorax. This flap was not removed but just reflected anteriorly to reveal the metathoracic ganglion. The lesions in separate locusts were: cutting the left metathoracic nerve root 5; cutting all nerve roots on the left side of the metathoracic ganglion; and removing the metathoracic ganglion completely. The flap of cuticle was folded back into position and the edges of the wound sealed with molten wax to prevent loss of haemolymph. All animals survived without any obvious distress (some of them were tested the following day) though with clear disabilities confirming the lesions.

### Data collection

We collected data during entry to chill coma from 93 intact locusts and 79 semi-intact preparations. To correlate CO_2_ release with bursts of electrical activity recorded from the head or thorax we performed experiments on 55 locusts. The early experiments established that recording from wing muscles was the most reliable approach, as has been found previously ^11^,^12^, and the quantitative comparisons were made using the last 20 locusts. Whole animal electromyographic recordings from wing and leg muscles and without respirometry were made using another 38 locusts. Measurements from all 38 were used to quantify temperature. Of the 38, 2 were not lesioned, 12 had the left metathoracic N5 cut, 16 had all metathoracic nerve roots on the left side cut and 8 had the metathoracic ganglion completely removed.

Initial characterization of the negative DC shift of field potential that is a signature of SD used 12 locusts and [K^+^]o during chilling was measured in another 6 locusts. The spreading nature of SD was confirmed by recording FP at two different locations in the mesothoracic ganglion in 7 locusts. In 14 locusts FP was recorded simultaneously with neural activity in the left metathoracic N5 (cut just distal to the electrode). FP was recorded simultaneously with extracellular recording of the DCMD axon in the connective between the meso- and metathoracic ganglia in 18 locusts and with intracellular DCMD axon recording in 10 locusts. During these recordings the DCMD was visually stimulated by moving objects towards the head in the visual hemi-field contralateral to the recorded axon. Intracellular recordings from the neuropil segments of wing muscle motoneurons, identified with reference to DL emg activity during flight sequences evoked by wind stimulation of the head, were made from 12 locusts.

### Data analysis

Electrophysiological recordings were filtered during acquisition, as described above, but also during subsequent analysis using the tools available in Clampfit 10.3 (pClamp, Molecular Devices, Inc.). Different filter settings were used to filter out 60-cycle mains interference and high frequency noise. For display purposes, brief electrical artifacts associated with moving the saline inlet tube around the preparation were cut from the recordings by inserting straight line sections between cursors placed on either side of the artifact. These are so brief as to be undetectable in the figure traces. For traces of slow events (temperature, FP, [K^+^]o) that were reconstructed as graphs in Sigmaplot 12.5 (Systat Software, Inc.) the number of data points was reduced by decimation.

For the intact animal experiments, the thoracic temperature was measured at the middle of the two peaks of CO_2_ release and at the beginning of the first clear burst of electrical activity from a wing muscle. For the semi-intact preparations, the saline temperature adjacent to the mesothoracic ganglion was measured at the end of neural firing recorded either extracellularly or intracellularly and at the half-maximal amplitude of the negative DC shift of FP. Thus the intact animal experiments better characterized the start of events leading to CT_min_ whereas the semi-intact experiments better characterized the occurrence of CT_min_.

Statistical analysis was performed using the software in Sigmaplot 12.5. Data were tested for normality (Shapiro-Wilk test) and equal variance (Levene median test). Parametric data were compared using ANOVA followed by Holm-Sidak tests for multiple comparisons. Descriptive data are reported in the results text as mean ± standard deviation (SD). For comparison of means, parametric data are illustrated in the figures as mean ± standard error (SE). Non-parametric data are reported as median ± interquartile range (IQR) and statistically compared using Mann-Whitney rank sum tests. Statistically significant data (P < 0.05) are indicated by lettering (columns with the same letter are not statistically different). Statistics are reported in the Results text.

## References

1 Hazell, S.P. & Bale, J. S. Low temperature thresholds: are chill coma and CT(min) synonymous? J Insect Physiol 57, 1085–1089, doi:10.1016/j.jinsphys.2011.04.004 (2011).

2 Macmillan, H.A. & Sinclair, B. J. Mechanisms underlying insect chill-coma. J Insect Physiol 57, 12–20, doi:10.1016/j.jinsphys.2010.10.004 (2011).

3 Overgaard, J. & MacMillan, H. A. The integrative physiology of insect chill tolerance. Annu Rev Physiol 79, 187–208, doi:10.1146/annurev-physiol-022516-034142 (2017).

4 Andersen, J. L. et al. How to assess Drosophila cold tolerance: chill coma temperature and lower lethal temperature are the best predictors of cold distribution limits. Funct Ecol 29, 55–65, doi:10.1111/1365-2435.12310 (2015).

5 Findsen, A., Pedersen, T. H., Petersen, A. G., Nielsen, O.B. & Overgaard, J. Why do insects enter and recover from chill coma? Low temperature and high extracellular potassium compromise muscle function in Locusta migratoria. J Exp Biol 217, 1297–1306, doi:10.1242/jeb.098442 (2014).

6 Findsen, A., Overgaard, J. & Pedersen, T. H. Reduced L-type Ca2+ current and compromised excitability induce loss of skeletal muscle function during acute cooling in locust. Journal of Experimental Biology 219, 2340–2348, doi:10.1242/jeb.137604 (2016).

7 MacMillan, H. A., Findsen, A., Pedersen, T.H. & Overgaard, J. Cold-induced depolarization of insect muscle: differing roles of extracellular K+ during acute and chronic chilling. J Exp Biol 217, 2930–2938, doi:10.1242/jeb.107516 (2014).

8 Anderson, R.L. & Mutchmor, J. A. Temperature acclimation and its influence on electrical activity of nervous system in three species of cockroaches. Journal of Insect Physiology 14, 243-246 &, doi:Doi 10.1016/0022-1910(68)90034-6 (1968).

9 Staszak, D.J. & Mutchmor, J. A. Influence of temperature on chill-coma and electrical activity of central and peripheral nervous systems of american cockroach, Periplaneta americana. Comp. Biochem. Physiol. 45, 909–923, doi:Doi 10.1016/0300-9629(73)90327-7 (1973).

10 Bradfisch, G. A., Drewes, C.D. & Mutchmor, J. A. The effects of cooling on an identified reflex pathway in the cockroach (Periplaneta americana), in relation to chill-coma. J. Exp. Biol. 96, 131–141 (1982).

11 Goller, F. & Esch, H. Comparative study of chill-coma temperatures and muscle potentials in insect flight muscles. Journal of Experimental Biology 150, 221–231 (1990).

12 Hosler, J. S., Burns, J.E. & Esch, H. E. Flight muscle resting potential and species-specific differences in chill-coma. J Insect Physiol 46, 621–627 (2000).

13 Esch, H. The effects of temperature on flight-muscle potentials in honeybees and cuculiinid winter moths. Journal of Experimental Biology 135, 109–117 (1988).

14 Andersen, J. L., MacMillan, H.A. & Overgaard, J. Muscle membrane potential and insect chill coma. J Exp Biol 218, 2492–2495, doi:10.1242/jeb.123760 (2015).

15 Rodgers, C. I., Armstrong, G. A. B. & Robertson, R. M. Coma in response to environmental stress in the locust: A model for cortical spreading depression. Journal of Insect Physiology 56, 980–990, doi:10.1016/j.jinsphys.2010.03.030 (2010).

16 Armstrong, G. A., Rodriguez, E.C. & Meldrum Robertson, R. Cold hardening modulates K+ homeostasis in the brain of Drosophila melanogaster during chill coma. J Insect Physiol 58, 1511–1516, doi:10.1016/j.jinsphys.2012.09.006 (2012).

17 Leao, A. A. P. Spreading depression of activity in the cerebral cortex. Journal of Neurophysiology 7, 359–390 (1944).

18 Dreier, J.P. & Reiffurth, C. The stroke-migraine depolarization continuum. Neuron 86, 902–922, doi:10.1016/j.neuron.2015.04.004 (2015).

19 Rodgers, C. I. et al. Stress preconditioning of spreading depression in the locust CNS. Plos One 2, doi:ARTN e13661 0.1371/journal.pone.0001366 (2007).

20 Armstrong, G. A. B. et al. Glial Hsp70 Protects K+ Homeostasis in the Drosophila Brain during Repetitive Anoxic Depolarization. Plos One 6, doi:10.1371/journal.pone.0028994 (2011).

21 Spong, K. E., Rodriguez, E.C. & Robertson, R. M. Spreading depolarization in the brain of Drosophila is induced by inhibition of the Na+/K+-ATPase and mitigated by a decrease in activity of protein kinase G. J Neurophysiol 116, 1152–1160, doi:10.1152/jn.00353.2016 (2016).

22 Spong, K. E., Andrew, R.D. & Robertson, R. M. Mechanisms of spreading depolarization in vertebrate and insect central nervous systems. J Neurophysiol 116, 1117–1127, doi:10.1152/jn.00352.2016 (2016).

23 Pietrobon, D. & Moskowitz, M. A. Chaos and commotion in the wake of cortical spreading depression and spreading depolarizations. Nat Rev Neurosci 15, 379–393, doi:10.1038/nrn3770 (2014).

24 Spong, K. E., Mazzetti, T.R. & Robertson, R. M. Activity dependence of spreading depression in the locust CNS. Journal of Experimental Biology 219, 626–630, doi:10.1242/jeb.132456 (2016).

25 Bogdanov, V. B. et al. Susceptibility of primary sensory cortex to spreading depolarizations. J Neurosci 36, 4733–4743, doi:10.1523/JNEUROSCI.3694-15.2016 (2016).

26 von Bornstadt, D. et al. Supply-demand mismatch transients in susceptible peri-infarct hot zones explain the origins of spreading injury depolarizations. Neuron 85, 1117–1131, doi:10.1016/j.neuron.2015.02.007 (2015).

27 Macmillan, H. A., Williams, C. M., Staples, J.F. & Sinclair, B. J. Metabolism and energy supply below the critical thermal _min_imum of a chill-susceptible insect. J Exp Biol 215, 1366–1372, doi:10.1242/jeb.066381 (2012).

28 Sinclair, B. J., Klok, C.J. & Chown, S. L. Metabolism of the sub-Antarctic caterpillar Pringleophaga marioni during cooling, freezing and thawing. Journal of Experimental Biology 207, 1287–1294, doi:DOI 10.1242/jeb.00880 (2004).

29 Stevens, M. M., Jackson, S., Bester, S. A., Terblanche, J.S. & Chown, S. L. Oxygen limitation and thermal tolerance in two terrestrial arthropod species. J Exp Biol 213, 2209–2218, doi:10.1242/jeb.040170 (2010).

30 Lighton, J. R. B. & Turner, R. J. Thermolimit respirometry: an objective assessment of critical thermal maxima in two sympatric desert harvester ants, Pogonomyrmex rugosus and P-californicus. Journal of Experimental Biology 207, 1903–1913, doi:10.1242/jeb.00970 (2004).

31 Groenewald, B., Hetz, S. K., Chown, S.L. & Terblanche, J. S. Respiratory dynamics of discontinuous gas exchange in the tracheal system of the desert locust, Schistocerca gregaria. Journal of Experimental Biology 215, 2301–2307, doi:10.1242/jeb.070995 (2012).

32 Berman, T. S., Ayali, A. & Gefen, E. Neural control of gas exchange patterns in insects: locust density-dependent phases as a test case. Plos One 8, doi:ARTN e59967 10.1371/journal.pone.0059967 (2013).

33 Money, T. G. A., Sproule, M. K. J., Hamour, A.F. & Robertson, R. M. Reduction in neural performance following recovery from anoxic stress Is mimicked by AMPK pathway activation. Plos One 9, doi:10.1371/journal.pone.0088570 (2014).

34 Gray, J., Lee, J. & Robertson, R. Activity of descending contralateral movement detector neurons and collision avoidance behaviour in response to head-on visual stimuli in locusts. J Comp Physiol A 187, 115–129 (2001).

35 Kovac, H., Stabentheiner, A., Hetz, S. K., Petz, M. & Crailsheim, K. Respiration of resting honeybees. Journal of Insect Physiology 53, 1250–1261, doi:10.1016/j.jinsphys.2007.06.019 (2007).

36 Stoney, S. D., Jr. & Machne, X. Mechanisms of accommodation in different types of frog neurons. J Gen Physiol 53, 248–262 (1969).

37 Armstrong, G. A. B., Rodgers, C. I., Money, T. G. A. & Robertson, R. M. Suppression of spreading depression-like events in locusts by inhibition of the NO/cGMP/PKG pathway. J. Neurosci. 29, 8225–8235, doi:10.1523/JNEUROSCI.1652-09.2009 (2009).

38 Spong, K.E. & Robertson, R. M. Pharmacological blockade of gap junctions induces repetitive surging of extracellular potassium within the locust CNS. Journal of Insect Physiology 59, 1031–1040, doi:10.1016/j.jinsphys.2013.07.007 (2013).

39 Brisson, C. D., Hsieh, Y. T., Kim, D., Jin, A. Y. & Andrew, R. D. Brainstem neurons survive the identical ischemic stress that kills higher neurons: insight to the persistent vegetative state. PLoS One 9, e96585, doi:10.1371/journal.pone.0096585 (2014).

40 Robertson, R. M. Thermal stress and neural function: adaptive mechanisms in insect model systems. Journal of Thermal Biology 29, 351–358, doi:10.1016/j.jtherbio.2004.08.073 (2004).

41 Robertson, R.M. & Money, T. G. A. Temperature and neuronal circuit function: compensation, tuning and tolerance. Current Opinion in Neurobiology 22, 724–734, doi:10.1016/j.conb.2012.01.008 (2012).

42 Andersen, M. K., Jensen, S.O. & Overgaard, J. Physiological correlates of chill susceptibility in Lepidoptera. J Insect Physiol 98, 317–326, doi:10.1016/j.jinsphys.2017.02.002 (2017).

43 Robertson, R.M. & Pearson, K. G. A preparation for the intracellular analysis of neuronal activity during flight in the locust. J. Comp. Physiol. A 146, 311–320 (1982).

